# Real-time predictions of reservoir size and rebound time during antiretroviral therapy interruption trials for HIV

**DOI:** 10.1101/038091

**Authors:** Alison L. Hill, Daniel I.S. Rosenbloom, Edward Goldstein, Emily Hanhauser, Daniel R. Kuritzkes, Robert F. Siliciano, Timothy J. Henrich

## Abstract

Monitoring the efficacy of novel reservoir-reducing treatments for HIV is challenging. The limited ability to sample and quantify latent infection means that supervised antiretroviral therapy (ART) interruption studies are generally required. Here we introduce a set of mathematical and statistical modeling tools to aid in the design and interpretation of ART-interruption trials. We show how the likely size of the remaining reservoir can be updated in real-time as patients continue off treatment, by combining the output of laboratory assays with insights from models of reservoir dynamics and rebound. We design an optimal schedule for viral load sampling during interruption, whereby the frequency of follow-up can be decreased as patients continue off ART without rebound. While this scheme can minimize costs when the chance of rebound between visits is low, we find that the reservoir will be almost completely reseeded before rebound is detected unless sampling occurs at least every two weeks and the most sensitive viral load assays are used. We use simulated data to predict the clinical trial size needed to estimate treatment effects in the face of highly variable patient outcomes and imperfect reservoir assays. Our findings suggest that large numbers of patients – between 40 and 150 – will be necessary to reliably estimate the reservoir-reducing potential of a new therapy and to compare this across interventions. As an example, we apply these methods to the two “Boston patients”, recipients of allogeneic hematopoietic stem cell transplants who experienced large reductions in latent infection and underwent ART-interruption. We argue that the timing of viral rebound was not particularly surprising given the information available before treatment cessation. Additionally, we show how other clinical data can be used to estimate the relative contribution that remaining HIV+ cells in the recipient versus newly infected cells from the donor made to the residual reservoir that eventually caused rebound. Together, these tools will aid HIV researchers in the evaluating new potentially-curative strategies that target the latent reservoir.

## Introduction

Twenty years after the introduction of combination antiretroviral therapy (cART) for HIV infection, the search continues for a cure, an intervention that would allow infected individuals to discontinue all treatments without experiencing viral rebound. One promising approach to achieve a cure is to eradicate latent virus that remains in resting CD4^+^ T cells despite long-term cART [1]. Pharmacologic agents that reactivate viral gene expression, collectively called latency-reversing agents (LRA), are undergoing preliminary clinical evaluation [2–4]. A major unknown regarding the potential efficacy and safety of LRAs is how much the latent reservoir (LR) must be reduced in order to delay or prevent viral rebound following cART interruption. Answering this question is an important prerequisite for initiating or scaling up clinical trials. For example, should a compound that reactivates 90% of latently infected cells *in vitro* be moved into clinical trials, or would this compound provide insufficient benefit relative to the risk of interrupting cART? What if it’s 99%?

Case reports of patients who achieved reduction in the latent reservoir by different means give some guidance. A greater than 3.5 log-reduction in the “Berlin patient,” in the setting of the Δ 32 CCR5 mutation, has lead to a cART-free period without viral rebound that is now 8 years in length [5]. At least 2 log-reductions in two hematopoietic stem-cell transplant recipients, the “Boston patients,” resulted in cART-free remissions of 3 and 8 months before viral rebound [6, 7]. An early-treated neonate, the “Mississippi baby,” achieved an LR size at least 2.5 logs smaller than the typical adult size and experienced rebound after 27 months [8]. An early treatment initiation case with LR size ≈ 3 logs below the typical size rebounded after 50 days [9].

We recently developed a mechanistic mathematical model to predict the time to rebound following reservoir reduction [10]. This model describes the relationship between reservoir size and time to viral rebound, and it predicts large inter-patient variability in response to identical treatment regimes. It also suggests that rebound after many months or even years of remission is a likely outcome, thereby independently predicting the observations of recent case studies [7, 8].

Despite the insight provided by these disparate case studies and mathematical modeling, there are many open questions about the design and interpretation of trials to assess interventions that reduce the pool of latent virus in HIV-infected individuals. One major challenge is the limited dynamic range of assays that measure the size of the latent reservoir [11–14]. Consequently, we often do not know the reservoir-reduction induced by the treatment, limiting our ability to predict outcomes based on previously observed or modeled data.

Absent direct measurement, can we estimate the LR size (and reduction efficacy) immediately following LRA therapy, based on the mechanism of action of the therapy or other biomarkers? Can we refine this estimate by observing long-term clinical outcomes of LRA therapy, in the face of highly variable patient outcomes? Another obstacle is the multi-year follow-up required to confirm that a patient is cured. How often and for how long should viral load measurements be taken during interruption trials, and can the likelihood of eventual rebound be updated in real-time as patients continue off treatment?

Many of these uncertainties were highlighted in the recent case of the “Boston patients” [6, 7]. Two HIV^+^ individuals with lymphoma received allogeneic hematopoietic stem cell transplants (HSCT), after which HIV viral RNA became undetectable by standard clinical assays. Both patients discontinued cART, and months later they remained HIV-free and were widely believed to have achieved long-term antiviral-free remission. However, after three and eight months, both patients experienced spontaneous and rapid viral rebound, suggesting that similar difficulties are likely to be encountered in future treatment interruption trials. Despite their impracticality for scale-up, HSCT studies are informative to HIV cure research, because they offer a model system to test the hypothesis that purging the latent reservoir can prevent rebound of the infection after interrupting cART. Understanding the likelihood of rebound and the mechanism of viral persistence in transplant studies is therefore a key facet of HIV cure research.

Here we present new quantitative approaches for understanding the dynamics of treatment interruption following reservoir reduction. We extend our mathematical modeling framework [10] to address the questions raised above, showing how this framework can assist with designing and interpreting outcomes of clinical trials for LRAs. We use clinical and virological data from the Boston patients as a case study to demonstrate the utility of these methods.

## Results

### Summary of the model of reservoir dynamics and rebound

Our basic model considers what happens *after* an LR-reducing therapy is administered and combination antiretroviral therapy (cART) is interrupted (Figure 1a). It is explained and analyzed in detail in a separate paper [10]. Briefly, we use a fully stochastic model of infection dynamics to calculate the probability of achieving a cure or, failing that, the duration of successful treatment interruption before viral rebound occurs, both as functions of the latent reservoir size following an intervention designed to reduce it. For the purpose of the model, this intervention may be a compound or strategy that reactivates and kills latently infected cells, an allogeneic hematopoietic stem cell transplant, or any other method of removing latently infected cells. The model tracks two CD4+ T cell types: productively infected activated cells, and latently infected resting cells. Latently infected cells can activate, die, or proliferate. Actively infected cells can produce a burst of virions, resulting in the infection of other cells, or die without producing virions. The model is designed to describe only the initial stages of rebound, where viral loads are well below typical setpoints.

**Figure 1.**
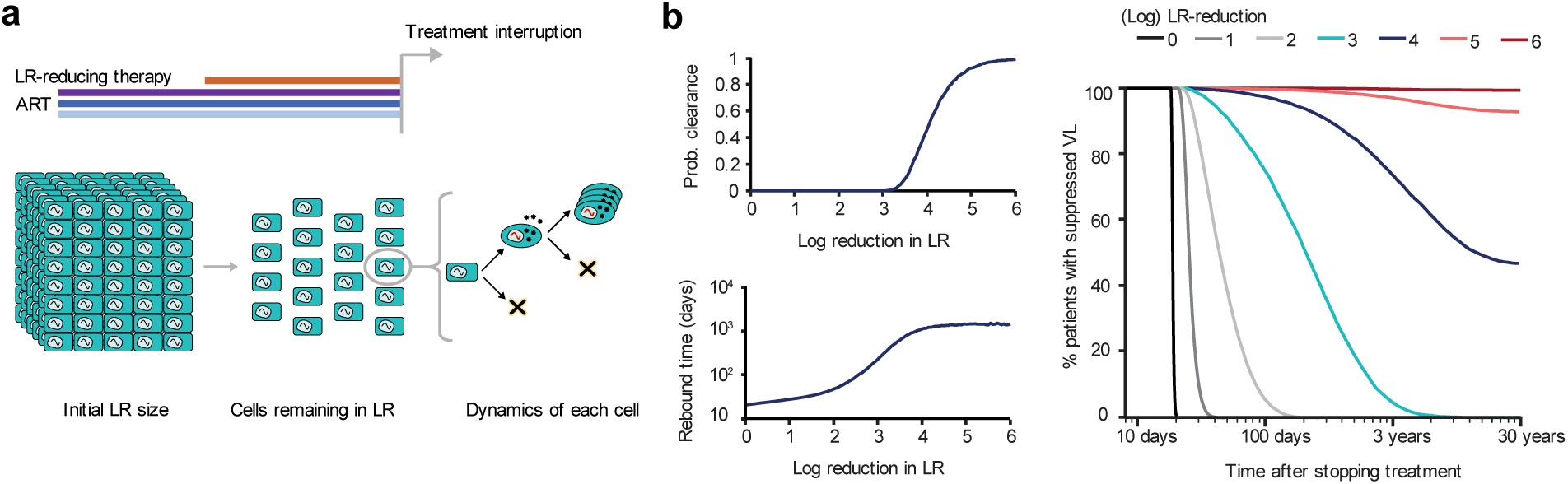
Summary of model of reservoir dynamics and rebound. a) Patients on fully suppressed ART are given an additional intervention to reduce the LR size. The stochastic model of viral dynamics following ART-interruption tracks both latently infected resting CD4+ T cells (rectangles) and productively infected CD4^+^ T cells (ovals). Viral rebound occurs if at least one remaining latently infected cell survives long enough to activate and produce a chain of infection events leading to detectable viremia. b) Model predictions for probability that the LR is cleared. Clearance occurs if all cells in the LR die before a reactivating lineage leads to viral rebound. c) Predicted median viral rebound times among patients who do not clear the infection. d) Survival curves (Kaplan-Meier plots) show the percentage of patients predicted to have not yet experienced viral rebound, as a function of the time after treatment interruption.

We concluded that model outcomes depend on a small number of parameters, which could be estimated from existing clinical data [10]. Specifically, the half-life of the pool of latently infected cells is estimated from studies of LR decay [15, 16], and the activation rate of latent cells and the viral growth rate can both be estimated from studies of supervised cART interruption [17, 18]. A fourth parameter, the probability that a single activated cell manages to establish a growing infection, is less certain and is the subject of both *in vitro* [19, 20] and population-genetic research [21–23] (reviewed in [24]).

The model has been used to estimate the relationship between the reservoir size and the possible outcomes following cART interruption. Here, we measure the reservoir size as the reduction from to the baseline value of a typical patient, i.e. around 1 infectious unit per million (IUPM) cells [12, 16]. We call this the *reduction efficacy.* If a patient begins LRA treatment with a higher or lower reservoir size than this, the reduction efficacy required to achieve the same outcome would be correspondingly higher or lower by the same amount. Our analysis is also relevant to patients in whom initial reservoir seeding was limited, such as during early treatment initiation. In this case, the reduction efficacy is simply the size of the reservoir relative to that of the aforementioned “typical” patient (who began cART during chronic infection).

The best outcome of reservoir-reduction, barring complete eradication, is that so few latently infected cells survive that none successfully reactivate and restart the infection. In this case, the infection has essentially been cured. If therapy fails to clear the infection, the next-best outcome is extension of the time until rebound. The results suggested three important findings. Firstly, reductions of 2 to 3 logs are required to delay rebound for a few months, while > 4-log reductions may be required for cure in most patients (Figure 1b). Secondly, patients may rebound even after experiencing several years off treatment without detectable viremia. These late rebounds pose a challenge for patient management. Thirdly, large inter-patient variation in times to rebound is expected, meaning that it is very difficult to predict outcomes for any individual patient (Figure 1c). This previous work, however, does not address the model’s ability to inform and interpret clinical trials. Here we show, using the “Boston patients” as case studies, how this model can be used to provide clinical guidance.

### Recovering reduction efficacy and LR size based on time of rebound

If the amount by which the LR is reduced is known exactly, then it is possible to predict time to rebound (Figure 1b). Yet measuring reduction efficacy directly from differences in frequency of latently infected cells observed before and after therapy may not be practical, due to the limited dynamic range of current assays and the difficulties in sampling large numbers of T cells from patients. Here we provide an approach to obtain estimates of reduction efficacy by supplementing the information from latency assays with the information gained by observing a patient’s rebound time (or absence of rebound). Intuition suggests that the longer a patient has been off treatment without rebound, the smaller the remaining reservoir size is likely to be. In other words, even if an investigator has a rough initial estimate of the reduction efficacy of an intervention (based on latency assays), this estimate continually should be revised upward with each subsequent negative viral load assay over the course of the interruption trial. Formally, we can use Bayesian updating to estimate the post-reduction reservoir size.

The Bayesian approach to estimate reservoir size requires two inputs: a distribution describing our knowledge about the reservoir size *before* the treatment interruption, known as the *prior* distribution, and a distribution describing how consistent a given reservoir size is with the observed rebound time (or the observed period of time without rebound), called the *likelihood.* The prior can be constructed based on any data available prior to observing the period of treatment interruption, such as viral outgrowth or PCR-based assays of the latent reservoir [12, 13], while the likelihood comes from the distribution of rebound times predicted by the model (Figure 1). The product of the prior and the likelihood, when normalized by a constant factor, gives the *posterior* distribution, reporting how probable each possible reservoir size is after accounting for all information. By this method, the observed clinical follow-up serves to narrow down the (often very broad) reservoir size estimates obtained from other means.

Illustrative estimates using this approach are shown in Figure 2. Calculation details are provided in the Methods. The prior distribution (Figure 2a) is constructed by assuming that immediately prior to cART-interruption, the reservoir was sampled at a level of 100 times the pre-reduction frequency of latently infected cells. For example, if there was 1 infectious unit per million cells (IUPM) before the reservoir was reduced, then we suppose that 100 million cells were sampled after reduction, none of which tested positive. To compute a prior from this negative assay result, we used a method based on Poisson sampling of the reservoir (summarized in Methods and detailed in [14]). This prior represents a conservative “rebuttable presumption” that reductions near 100-fold were the most likely outcome of therapy, and that a substantially greater reductions, while possible, should only be believed in the presence of convincing evidence (e.g., very long period of treatment interruption without rebound). This behavior is illustrated by the gradual slope downward on the right side of the graph in Figure 2a. The prior also treats reduction substantially less than 100-fold as very unlikely, as otherwise the post-treatment sample would almost certainly have returned a positive result; this behavior is illustrated by the sharp plummeting to the left on the log y-axis. Figure 2b shows the posterior estimates of the LR reduction efficacy, taking into account the observed time of rebound. A point estimate can be determined by taking the median of the posterior distribution, and 95% credible intervals can be constructed by taking the reduction efficacies between which 95% of the distribution falls. The same method can be used to estimate reservoir sizes during the course of trial, knowing only that rebound has *not* occurred prior to a certain time, but might occur at some point in the future (Figure 2c).

**Figure 2.**
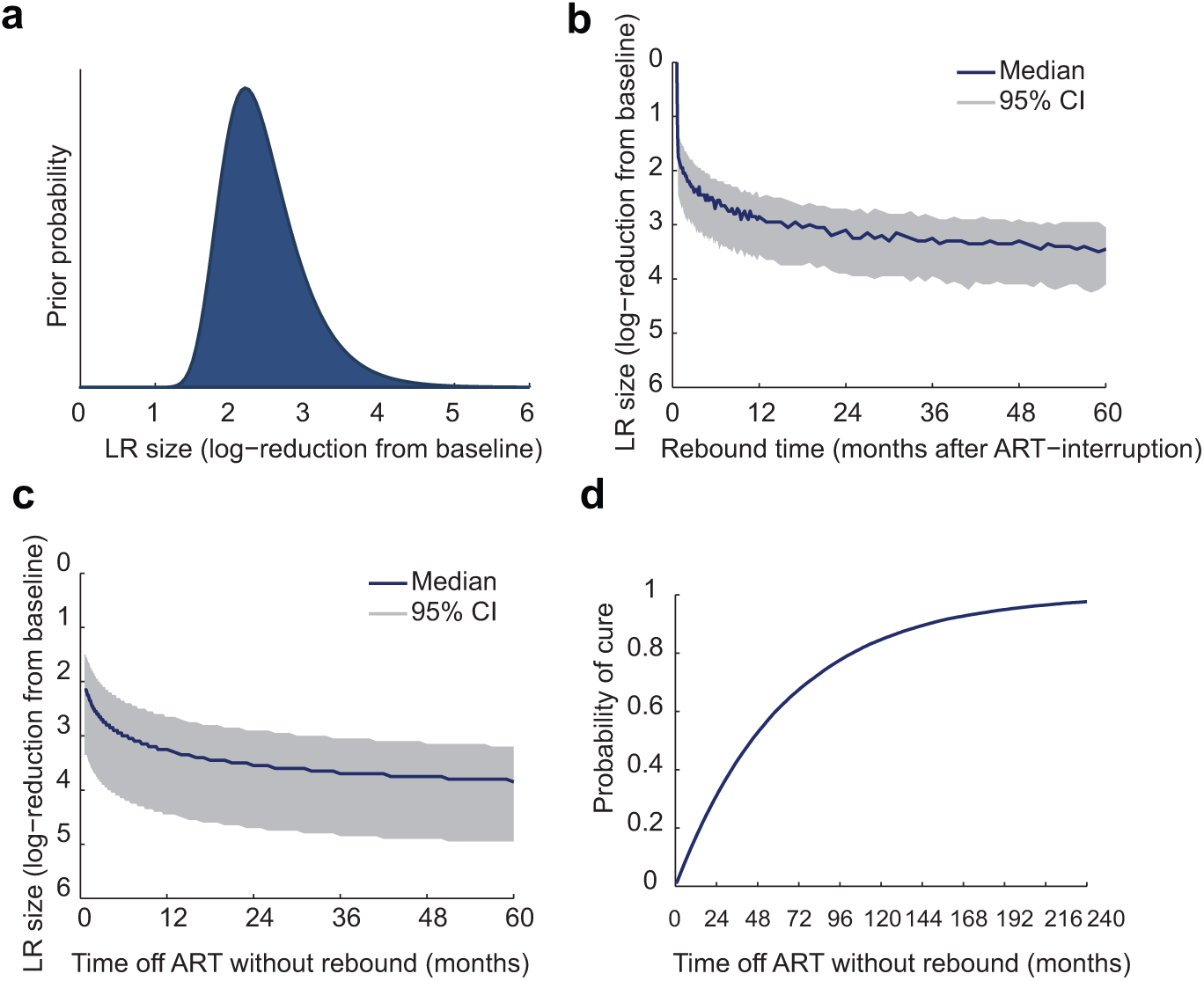
Predicting and interpreting the outcomes of treatment interruption when the reservoir reduction is unknown. A Bayesian approach is used to integrate information from reservoir assays with model predictions to relate the time absence or occurrence of viral rebound following ART-interruption to the remaining reservoir size. a) Prior distribution for the (unknown) reservoir reduction. We use as a prior the post-test likelihood for each reservoir size after a negative viral outgrowth assay (see Methods). b) The posterior probability for the LR reduction, given that *rebound has occurred,* using the posterior median (and 95% credible interval). c)The posterior probability for the LR reduction, as a function of the current time off treatment *without* rebound, using the posterior median (and 95% credible interval). d) The probability of ultimate reservoir clearance (cure) as a function of the current time off treatment *without* rebound. The initial cure probability is near zero and again takes over a decade to reach high values. Note that the reservoir reduction may either refer to the decrease in the number of latently infected cells in a given patient after administering a latency-reducing therapy, or, the factor by which initial reservoir seeding was limited, relative to a typical chronically infected patient.

Using this approach, 3 months without rebound implies an LR reduction of 500-fold (95% range 110 – 7,900-fold), while rebound *at* 3 months would suggest a 250-fold reduction (95% range 70 – 1,200-fold). For observations at 2 years, the corresponding estimates without rebound are 3,500-fold [250 – 50,000] and with rebound are 1,300-fold [500 – 7,900]. Note that observing rebound, as opposed to the absence of rebound, is predicted to lead to lower estimates for LR-reduction (higher estimates for size) and narrows the confidence interval on these estimates. However, all these results illustrate how the high variability between patient outcomes makes it difficult to estimate precise estimates of reservoir size reduction.

### Adaptive probability of cure

Intuition suggests that the longer a patient has been off treatment without rebound, the more likely it is that they will never experience rebound, thereby achieving a cure. As above, we can formalize this intuition using a Bayesian approach. Since the model gives a probability of cure for each possible reservoir size (Figure 1b), we simply estimate a patient’s cure probability by computing a weighted sum over that patient’s posterior distribution of LR sizes. Since this posterior distribution changes over time (Figure 2c), the expected cure probability likewise changes over time (Figure 2d), increasing with the duration of the interruption. Assuming, as above, that a post-reduction sample of 100 million cells test negative for latent infection, the cure probability immediately following reduction is only around 1% and takes over 10 years to reach > 90%.

### Required sample size to power studies of LR-reduction

If we consider a population of patients, then one major unknown is how many patients will be required in a given clinical trial to reliably estimate the reservoir reduction efficacy (i.e. to “power” the study). We can estimate this number using simulated patient data (details provided in the Methods). We assume that all patients respond identically to treatment, so that the probability that any cell will be removed from the LR (and hence the expected value of the LR-reduction) is the same for everyone. We define an adequately powered study size to be one for which at least 95% of trials of this size would result in 95% credible intervals for the reservoir-reduction that contain the true value and are less than 1 log wide. The calculation can be repeated for any desired definition of adequate study power. Figure 3 shows how many patients are required to achieve this goal. If the parameters governing viral dynamics are identical in all patients (best estimates from [10]), then trials as small as 5–15 individuals can reliably estimate reservoir reductions of up to 4 logs, while cohorts of 40 to 150 individuals are needed to resolve reductions greater than 4 logs. For large reductions, outcomes are more variable and rebound may even occur after the study follow-up period, which we assume here is 10 years.

**Figure 3.**
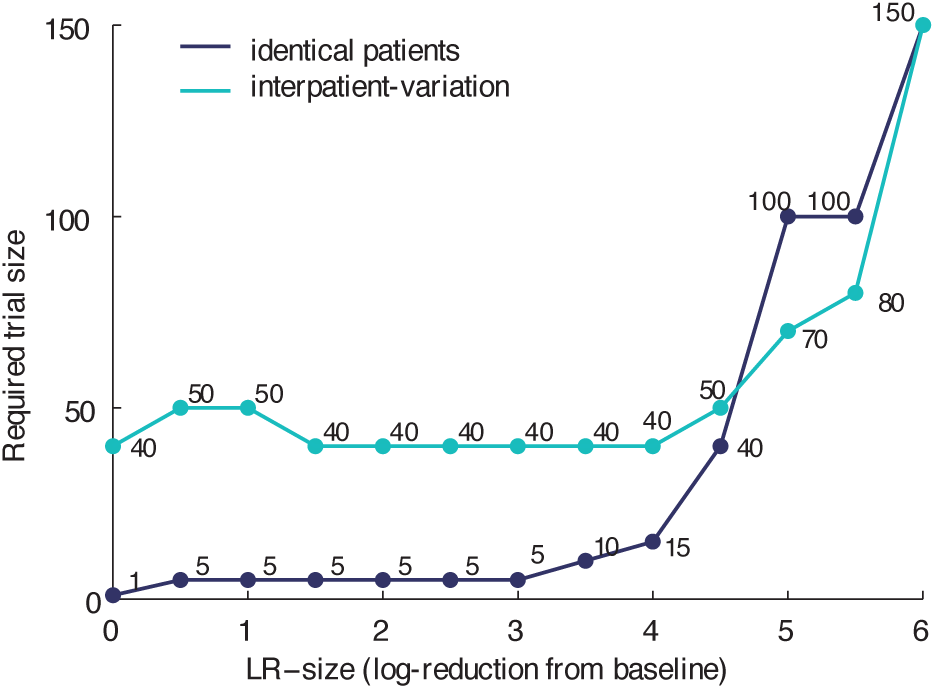
The required trial size to accurately and precisely estimate the LR-reduction. For each possible known LR-reduction (x-axis), we determine the number of patients in a trial (y-axis) needed so that at least 95% of trials of this size would result in 95% credible intervals for the estimated reduction that contain the true value and are less than 1 log wide. We sample hypothetical patients using our model, either with identical best-estimated parameter values (dark blue line), or allow interpatient variation from a range of possible values (cyan line); see distributions and data sources described in [10] and). We assume that all patients in the trial experience the same reservoir reduction (with only binomial variation in the actual number of cells remaining), and that patients are followed for a maximum of 10 years.

### Determining the optimal frequency of sampling during clinical trials

Our model results [10] as well as recent case studies [7, 25], have suggested that long-term patient follow-up is necessary for LR-reducing interventions, due to the risk of viral rebound even after long periods of remission. Frequent viral load testing for years after cART-interruption is expensive and time-consuming. Since rebound becomes less likely as a patient continues off therapy without viremia, it may be reasonable to decrease the frequency of viral load testing as time goes on. This model can be used to design an adaptive method for choosing the sampling intervals.

Efficient sampling can be achieved by choosing intervals in which the probability of rebound is constant, set to a pre-defined tolerance level. For example, we may decide that between each scheduled viral load test, we can tolerate that 5% of the suppressed study participants experience viral rebound. The Bayesian approach above can be used to calculate the fraction of suppressed patients expected to experience rebound, allowing us to choose sampling intervals meeting the desired risk tolerance. Again supposing that 100 million cells test negative for latent infection in an outgrowth assay, Figure 4a shows that, as expected, the recommended sampling frequency is initially high, and then drops off after months-to-years off treatment without rebound. Allowing for a 5% chance of failure between tests, the sampling frequency peaks at 4 times weekly and falls off to less than once per week after about 100 days. After about 1.5 years off treatment, the frequency drops to less than once per month. If we allow a 10% risk of failure between tests, sampling frequency peaks at only twice weekly and remains lower than in the 5% case. Importantly, the recommended testing frequency depends on prior information about LR reduction; if fewer than 100 million cells are sampled and tested for viral outgrowth following LR reduction, then recommended frequencies would increase.

**Figure 4.**
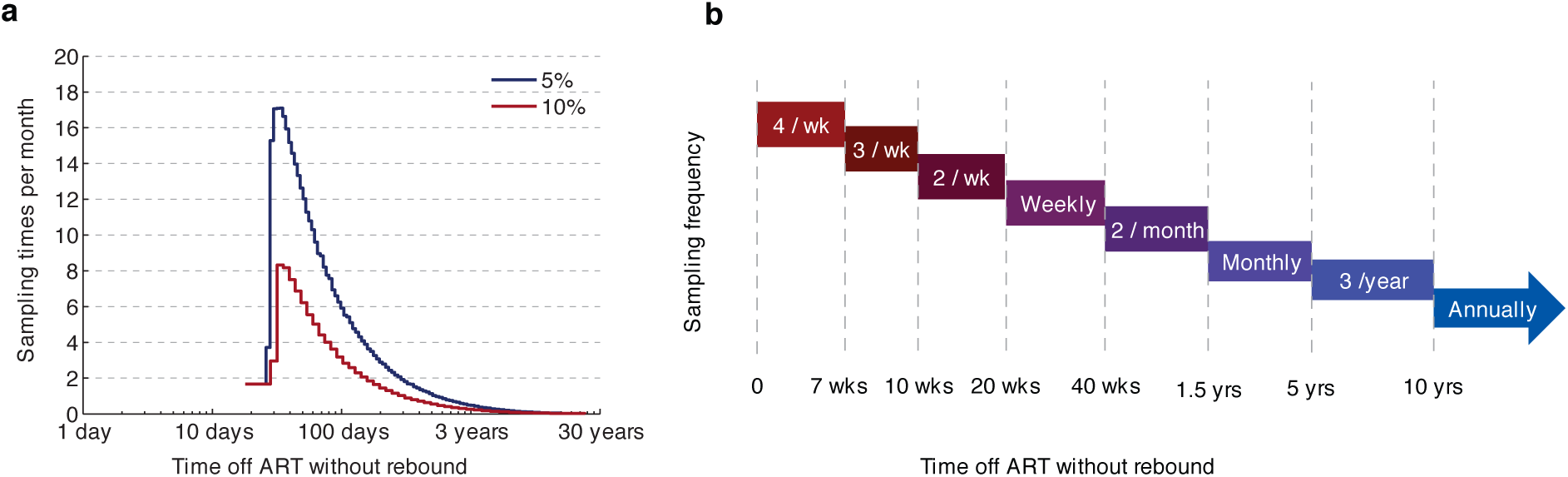
The optimal frequency of viral load sampling during supervised treatment interruptions. a) We calculate sampling times such that the probability of viral rebound between each test is equal and small - either 5% (dark blue line) or 10% (red line). These times are then transformed into intervals and expressed as the number of recommended samples per month. For these results we use the same prior distribution for the reservoir size as Figure 2a. The recommended frequency starts off low, before jumping to high values, because even without any reservoir-reduction, rebound rarely occurs within the first two weeks in patients who have been on suppressive cART. More frequent initial monitoring may be advisable if it is suspected that patients were not suppressed before interruption. b) A simplified sampling scheme that involves only regular intervals and assures less than 5% chance of failure between measures.

Note that because patients without any LR-reduction generally take a few weeks to rebound, the recommended sampling frequency is low initially and only peaks about three weeks after ART-cessation. However, other clinical concerns may favor beginning high-frequency testing immediately after interruption. For example, if a patient was not fully compliant with cART before interruption, viral replication may not have been fully suppressed, allowing viral rebound to occur very quickly. The sampling intervals in Figure 4b accounts for this concern and provides a simplified scheme for 5% rebound tolerance that may be more convenient for clinical protocols.

This sampling scheme assumes that the goal is to keep the chance of rebound constant and low between follow-up visits, not to stop rebound as soon as possible. If one wants to ensure that rebound is caught very quickly and high level viremia is prevented, then sampling must continue at a very high frequency indefinitely. Viral load can go from below 50 copies/ml to 10^4^ in around 2 weeks in patients with intact immune responses (exponential growth rate around 0.4/day), and perhaps in in as little as five days in those without HIV-specific immunity, as seen in the example of the Boston patients [7] discussed below.

### Potential for reservoir reseeding during treatment interruptions

One controversial aspect of treatment interruption trials is that if viral rebound occurs and is undetected for a long time, the reservoir may be reseeded, thereby undoing any benefit of the reservoir-reducing intervention. The extent of this reseeding is believed to depend on the total number of cells newly infected during the time that rebound is undetected, which in turn depends on the concentration of virus in the body (approximated by viral load in RNA copies per ml of plasma) and the abundance of cells available to be infected (approximated by the concentration of CD4^+^ T cells in cells per *μ*l of plasma). As detailed in, models suggest that the product of viral load and CD4^+^ count, integrated over time, should determine the number of new latently infected cells generated. This expectation was confirmed during the period of LR seeding that occurs in acute infection [26], and we use the empirical relationship observed in that paper to estimate LR reseeding after rebound. As a worst-case scenario, we assume that viral rebound begins prior to the most recent negative viral load test, such that viral load is just below the limit of detection at this time, and that it continues to grow according to a simple dynamic model until the infection is detected at the next sampling time and ART is re-initiated immediately. Figure 5a demonstrates that if patients are sampled infrequently, the LR could quickly return to a pre-treatment value (≈ 1 IUPM). More sensitive assays can reduce the potential for reservoir reseeding if sampling occurs at least every two weeks. Most reservoir reseeding occurs during peak viremia, and once viral load reaches a set-point, reservoir sizes increase much more slowly over time.

**Figure 5.**
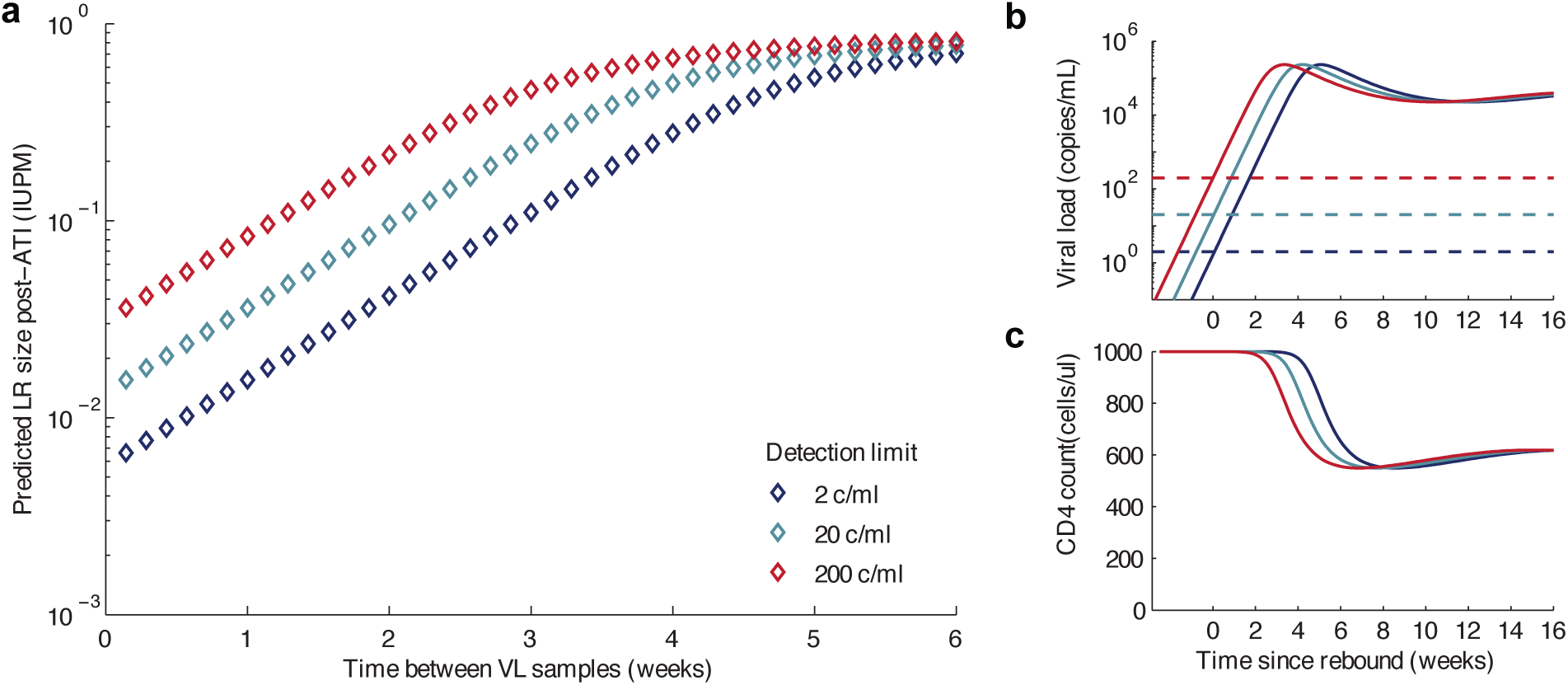
Impact of assay sensitivity and time between viral load samples on potential reservoir reseeding during rebound. We estimate how much re-seeding of the LR could occur between when viral rebound occurs and when it is detected, using a method for estimating LR sizes previously validated during acute infection [26]. We assume the LR size is very small (approx. zero) before interruption, though if it is larger these values represent the increase in size. LR size is measured as the frequency of infectious provirus among resting CD4^+^ T cells (IUPM = infectious units per million). We consider assay detection limits of 2 (dark blue), 20 (cyan), and 200 (red) copies/ml, and assume a worst-case scenario where viral load is just below this value at the last undetected sampling point. Smaller figures on the right show the viral load and CD4 count trajectories over time that generated the IUPM measurements in the larger figure. Time is measured relative to the time viral load reached the detection limit, and we assume detection does not occur until this value is surpassed.

### Sensitivity analysis for uncertainty in viral dynamic parameters

In the analyses presented thus far, we have assumed that the parameters governing reservoir dynamics and rebound – which are inputs to the model — are known and are identical between patients. Point estimates for each parameter come from multiple data sources and their derivation is detailed in [10]. However, uncertainty in these parameters remains. Here we consider how this uncertainty influences the methods we have described for interpreting rebound times. Details of the parameter ranges we considered, the numerical methods, and the results are provided in.

Including parameter uncertainty has two important effects. Firstly, it leads to more uncertainty to interpreting the reservoir size from time of rebound in individual patients. Delayed rebound could be due to few remaining latent cells or particularly favorable parameters, so longer remission times are less informative about the reduction efficacy alone. However, even with the broad parameter ranges considered, we find that these methods are still informative. For example, for the situation outlined in Figure 2, the pre-interruption median [95% CI] interval for the LR reduction based on laboratory assays alone was 2.16 [1.38, 3.41] logs. If the parameters are known precisely, then after 6 months without rebound this is increased to 3.00 [2.16, 4.21], while if the parameters are uncertain (with distributions given in, Table S1.1), then it is still increased but only to 2.63 [1.63, 4.03] (see, Figure S1.1). To test how much uncertainty could exist before rebound times become uninformative, we systematically increased the variance in one of the most important parameters governing rebound times - the rate at which latently infected cells become reactivated. We found that this information breakdown does not occur until the log-standard deviation increases by around four-fold (Figure S1.2). At this point, the observation of 6 months without rebound only decreases our estimate for the remaining reservoir size by around 0.1 log. This level of variation is unlikely to be biologically feasible, as it corresponds to around 10^7^-fold difference in the activation rate between the central 95% of individuals, suggesting our methods are robust to realistic levels of uncertainty.

Secondly, uncertainty in viral dynamics parameters means that the sample size required in a clinical trial to narrow down the efficacy of a reservoir-reducing intervention is increased (Figure 3, cyan line). To model this, we assumed that individual patients in a population have unique parameters which are randomly drawn from the ranges described (, [10]). In this case, study sizes of at least 40 are expected to be needed to reliably estimate any reduction efficacy. However, for nearly complete reservoir eradication, this interpatient variation actually decreases the required trial size to power studies. This occurs because when patients vary in reservoir reactivation dynamics, cure is predicted to be more common for small reservoir reductions but less likely for large reductions, due to the presence of a portion of patients with particularly favorable or unfavorable parameters. Since it is easier to estimate reduction efficacy from the occurrence of rebound than from the absence of it, fewer cures mean lower required trial sizes.

Overall, this sensitivity analysis demonstrates that even in the face of uncertainty about the viral dynamics parameters, observing rebound times always improves our estimates of the reservoir size following latency-reversing interventions beyond the estimates obtained from pre-interruption assays alone.

### Applications to allogeneic hematopoietic stem cell transplantation in the “Boston patients”

We consider two such HIV^+^ individuals who underwent cytotoxic chemotherapy and allogeneic hematopoietic stem cell transplantation to treat hematological malignancies. They have since become widely referred to as the “Boston patients.” Detailed clinical and virologic data for these individuals is reported elsewhere [6, 7], but summarized in Table 1, along with the results of our modeling work described in subsequent sections. Before transplant both patients had been on long-term cART, and had reservoir sizes (measured by presence of HIV DNA in PBMC) consistent with chronic infection. During transplant engraftment, transient viral load “blips” were detected although cART was continued. Three to four years post-transplant, engraftment appeared to be complete, as microchimerism assays detected residual host cells at levels of less than 0.001% in peripheral blood. Assays for plasma or latent virus returned no positive results. Due to lack of detectable HIV, both patients interrupted ART in a supervised manner. No free or integrated virus was detected until 3 and 8 months post-interruption, at which point plasma virus rapidly rebounded. Because these transplants occurred in the context of wildtype CCR5 (unlike the “Berlin patient” [5, 27]), the mechanism for delayed rebound is believed to be a reduction in the latent reservoir, suggesting it is well-suited to analysis by the modeling framework described in the previous section. We will examine how this analysis could have been used in real-time to interpret the outcomes of these patients, and how it can be used retrospectively. We will present and analyze the available clinical data on these patients in chronological order of how it was released to the public during the study. Note that for this analysis, we assume that the parameters governing reservoir and viral dynamics at the time of ART interruption are the same in these patients as in other treated HIV^+^ patients who did not receive transplants, including the rate of turnover and reactivation of latent cells, and the viral replication rate of reactivated lineages.

**Table 1.**
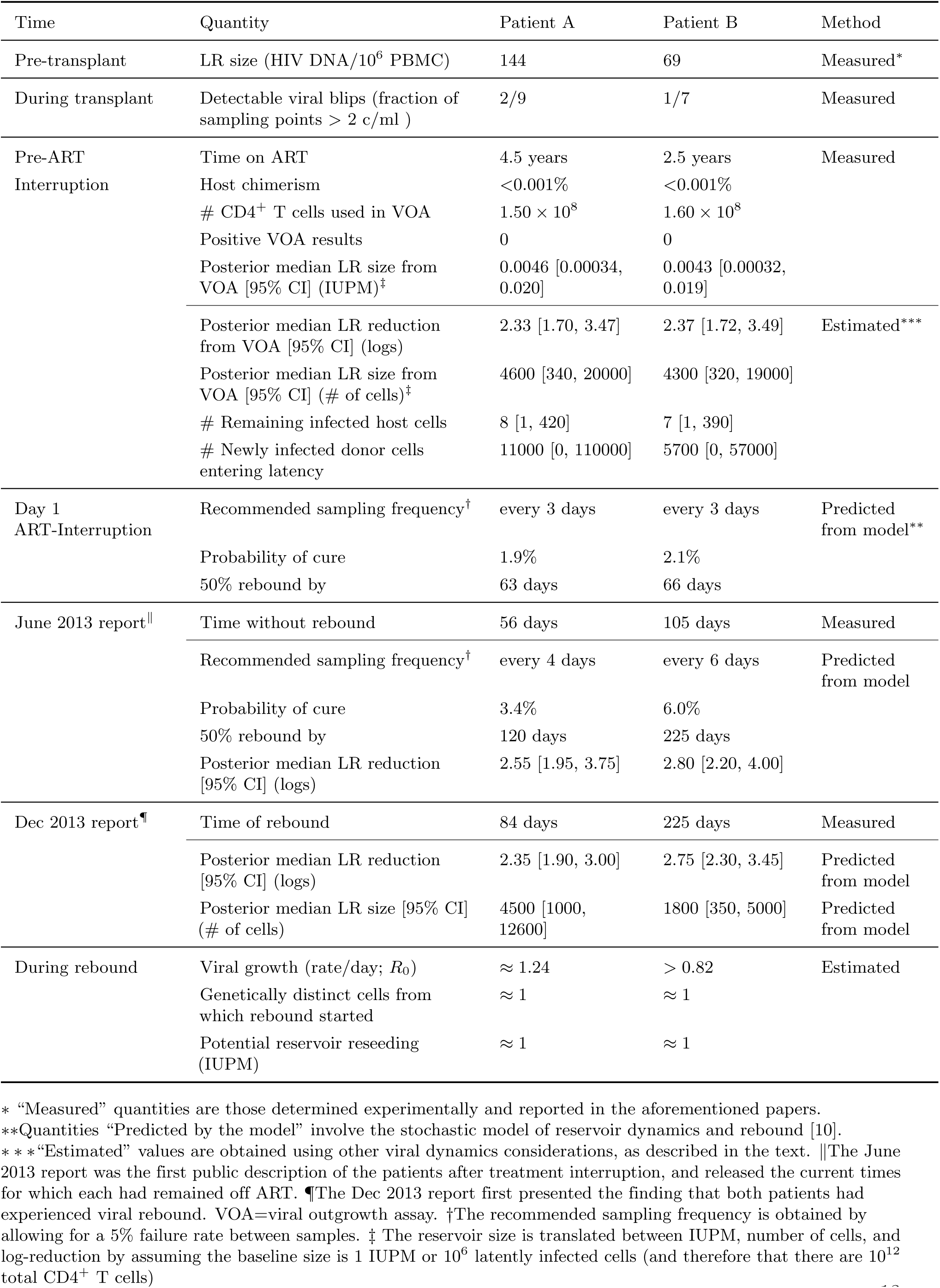
Reservoir quantification and rebound dynamics for the two “Boston patients”

### Predicting and interpreting outcomes for the Boston patients in real time

We examine the clinical and virologic data available for the “Boston patients” in the order released to the public, considering how additional modeling analysis could augment these data. At the time of treatment interruption, for each patient, a large number of CD4^+^ T cells were isolated and used in a viral outgrowth assay to measure the reservoir size. In each case the results were completely negative, and so using the approach described in the Methods and [14], we can create a post-test distribution for the likely size of reservoir. This distribution, described in terms of log-reduction from a baseline value of ≈ 1 IUPM, is created analogously to the distribution shown in Figure 2e. The median of this distribution is near 0.004 IUPM for both patients. Using this distribution as a prior for the unknown reservoir reduction, we can then estimate the probability each patient will be cured (i.e. not experience rebound), the day by which there is 50% chance that rebound will have occurred, and the recommended sampling frequency (reported in Table 1). Because it is still reasonably likely that the reservoir is quite large, the model predicts that sampling should occur frequently (every few days if the tolerance is 5% failure between samples), and that rebound is very likely to occur within the first few months.

The predictions can be updated throughout observation of the interruption trial. The first report of these patients was made public in June 2013 [28], describing that they had experienced antiretroviral-free remission for 8 and 15 weeks. Adding this information to the model results in more optimistic predictions (Table 1). The updated posterior median estimate for reservoir size is now reduced to 0.0028 and 0.0016 IUPM, for Patients A and B. This justifies only a small improvement in the expectation of cure, rising from 2% for each patient to 3% and 6%, respectively.

In December 2013 [29], it was reported that both patients had rebounded, after a total of 84 and 225 days. The posterior estimates for reservoir reduction based on this final piece of information are not much larger than the prior estimates obtained from the negative viral outgrowth assay results (median of 2.35 logs reduction versus 2.33 for Patient A, 2.75 versus 2.37 for Patient B, see Table 1), and the 95% credible intervals for the prior and the posterior overlap considerably. In other words, the final outcomes were not terribly surprising from the standpoint of the initial negative assay results, and had the same outgrowth assays been performed a second time prior to ART interruption (PBMC sample permitting), they may very well have shown a positive result.

### Estimating the size and source of the reservoir post-transplant

The case of the Boston patients highlights the fact that even after finding negative results in viral outgrowth assays of latency, a wide range of reservoir sizes are consistent with the data. Once rebound is observed, the time of rebound can help to narrow this range when interpreted in the context of a mathematical model of rebound dynamics. In the context of HSCT, observing rebound prompts the question of where the rebounding viral lineage originated. Two sources are possible: host cells that survived chemotherapy and graft-vs-host disease, and donor cells that later became infected and entered latency. A schematic of potential changes in reservoir size over the course of transplant, depicting both possible sources, is shown in Figure. Here we show how other experimental data can help us understand the source of rebound viremia following transplant and provide independent estimates for the size of the remaining reservoir. Note that in the case of the Boston patients, sequence analysis of pre- and post-transplant virus showed high relatedness, ruling out superinfection as a likely cause of viral rebound [6, 7].

**Figure 6.**
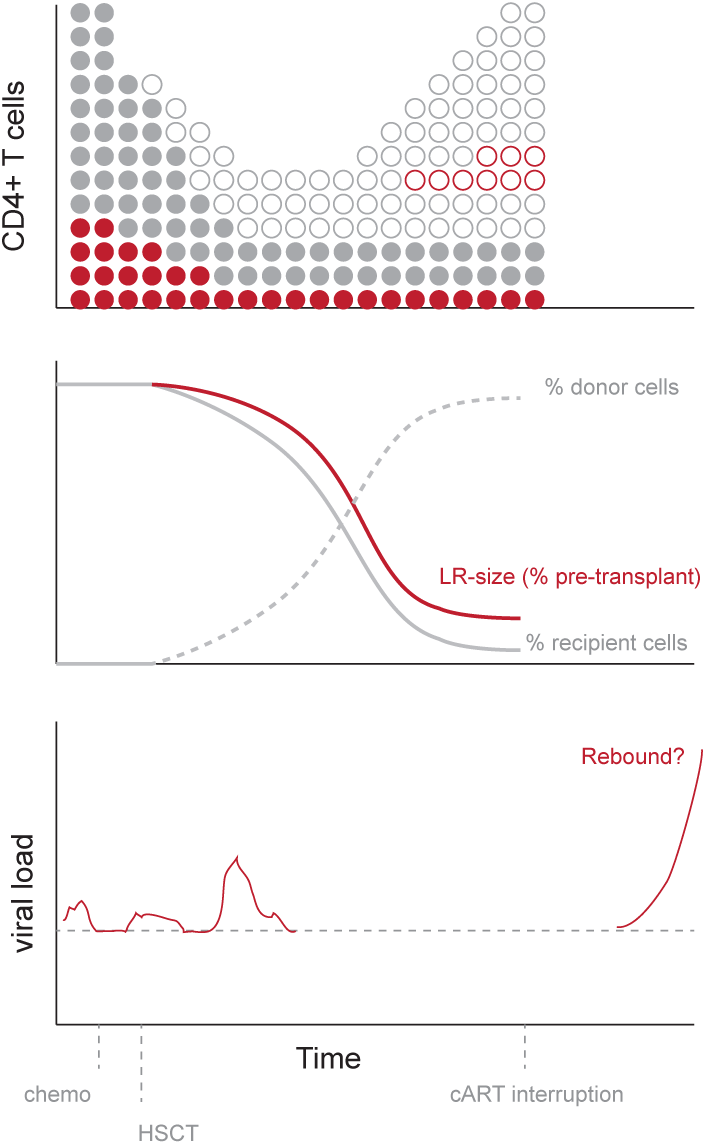
Schematic of potential cell and viral dynamics during hematopoietic stem cell transplant with suppressive cART. Solid circles: recipient cells. Open circles: donor cells. Red: HIV^+^ cells. The recipient patient starts out with high levels of CD4^+^ T cells, a small fraction of which are latently infected with HIV. Following conditioning chemotherapy, recipient cell levels drop. When donor cells are transplanted, recipient cells continue to decline as donor cells increase in number. If any ongoing viral replication occurs during engraftment, donor cells may become HIV-infected. Without new infections, the latent reservoir size should decrease proportionally to the frequency of recipient cells, but new infection of donor cells may quell this decrease. Viral blips may occur during transplant, perhaps due to imperfect cART adherence or immune-modulated viral re-activation. If cART is interrupted, then any remaining latently-infected cells – either from the recipient or donor – may reactivate and lead to viral rebound.

#### Remaining reservoir in host cells

Pre-transplant chemotherapy and post-transplant graft-vs-host-disease [6, 30, 31] may not be 100% effective at removing all host cells, and some host cells that comprised the LR may remain at a low frequency. The efficiency of engraftment in these patients was measured by allele-specific PCR of PBMC to identify host vs donor cells (“micro-chimerism”) [6], which found host cells to remain at a frequency less than 10^−5^ (0.001%) in the peripheral blood. This result suggests that the reservoir in host cells was reduced by at least 5 logs. To translate this value into an actual reservoir size (in terms of IUPM or number of infected cells), we need to know the pre-transplant LR size. The major difficulty in making this estimate is that reservoir sizes before transplant were only measured as the fraction of PBMC that contain HIV proviral DNA. Only a small portion of this integrated provirus is believed to be intact and able to produce virions [32]. Accordingly, the frequency of infectious cells as measured by viral outgrowth assays is much smaller than PBMC HIV DNA levels [12]. Moreover, there is only a weak correlation between the two measurements, and it is unclear which of these measurements is the best proxy for the true “functional” size of the reservoir from which viral rebound can occur. However, we can use them as upper and lower bounds on the reservoir size. As detailed in, we use previously determined ratios between these different assay measurements to estimate reservoir frequencies for Patient A and B of 7.5 (0.75, 420) x10^−6^ and 6.9 (0.69, 390) x10^−6^ respectively, which, assuming a total of 10^12^ resting CD4^+^ T cells, corresponds to 8 [1, 420] and 7 [1, 390] infected cells remaining.

#### New infection of donor cells

Both patients received transplants from HIV-negative donors, and so any possible contribution to the latent reservoir from donor cells must have been due to new latent infections occurring during the transplant procedure. Both patients also remained on cART during the entire engraftment procedure, which was thought to prevent any new infection. Several independent lines of evidence suggest that viral replication halts during optimal adherence to cART: treatment reduces viral replication several orders of magnitude *in vitro* [33, 34], evolution of the viral population appears to stop *in vivo* [35, 36], and intensified treatment does not further reduce viral load [37, 38]. Yet it is possible that short lapses in adherence [39], or the presence of cellular compartments with poor drug penetration [40], may compromise the effectiveness of cART, allowing low levels of new infection. Note that if this replication was ongoing until the time cART was interrupted, then viral rebound would be expected to occur immediately. The long delay until rebound implies that any ongoing replication was transient, so that upon treatment cessation rebound could not occur until a latent cell successfully reactivated. We estimated ranges for the number of latently infected donor cells formed by the time of cART-interruption using a method that was agnostic to the particular cause of ongoing replication. Following most viral dynamics models, we assumed that these cells are produced at a rate determined by the product of the viral load, the density of target CD4 cells, the infectivity, and the probability of infection resulting in latency. We used longitudinally observed levels of residual viremia and CD4 counts observed for each patient, and took bounds on the rates of infection and entry into latency during cART obtained from previous studies [26, 41]. As detailed in, the estimated ranges for the contribution to the latent reservoir from donor cells is 11,000 [0, 110,000] for Patient A and 5,700 [0, 57,000] for Patient B.

Despite the considerable uncertainty in these values and the estimated reservoir reductions, they do constrain the possible causes of viral rebound. Specifically, in Patient A, it is difficult to explain the observed rebound time without positing that some donor cells were newly infected. The 95% credible interval for the posterior estimate of the LR size estimated from the observed time of rebound suggests there were many more cells in the reservoir than were likely to remain from the recipients pre-transplant infection. However, if the efficiency of donor engraftment in tissues was lower than that in the peripheral blood, then enough latently infected host cells could have remained to explain the rebound time. In Patient B, on the other hand, the presence of new infection is not necessary to explain the observed rebound, as the upper bound on the number of residual host cells is consistent with the lower bound for the reservoir size estimated from the time of rebound.

### Analyzing rebound dynamics in the Boston patients

The dynamics of viral load following rebound can be used to estimate the rate of viral replication, to compare these two patients to other treatment interruption cohorts, and to examine potential reservoir re-seeding. For Patient A, there are two consecutive viral load values above the detection limit, before peak viremia and set-point are reached (900 c/ml at Day 84 and 130 000 c/ml at Day 88). From these, we estimate that the rate of exponential increase in viral load is *r* ≈ 1.24/day. This is higher than any of the values observed in non-transplant interruption studies (average 0.4/day, range 0.1-0.9) [10, 18, 42]. For Patient B, only one viral load value is available before cART was restarted, but from this (Day 225, 1 900 000 c/ml) and the last undetectable time point (Day 211, <20 c/ml), we can determine that *r* is at least 0.82/day. Overall these values suggest that viral replication is accelerated, or death of infected cells is inhibited, in HSCT patients due to loss of HIV-specific immune responses that may occur during the long period in which viral antigen is largely absent. In fact, these rates of viral increase are comparable to those observed during acute infection, prior to establishment of HIV-specific immune responses [43].

One of the predictions of the model of rebound dynamics is that if rebound occurs after a long delay, it is likely to be caused by a single cell that has reactivated from the latent reservoir. Significant delays in the rebound time only occur when the waiting time between latent cells reactivating is long, and since we assume that cells reactivate independently, the chance that more than one cell would start producing virus at nearly the same time is very low [10]. If this prediction is correct, then rebounding viremia should be highly genetically similar within the exponential growth phase, comparable to the viral population at the start of a single-founder HIV infection [44]. Phylogenetic analysis of rebounding viral lineages in both patients did indeed find that the sequences formed a distinct genetic cluster apart from the more diverse pre-transplant HIV DNA [7], thus supporting the mechanism of rebound suggested by the model. An alternate explanation for this lack of diversity in rebound viremia is that the latent reservoir is maintained by cellular proliferation, giving rise of clones of identical provirus [45–48]. In that case, multiple simultaneously reactivating cells may be indistinguishable from one another. A recent study has found, however, that most of these clones contain defective provirus [49], decreasing the plausibility of this alternate explanation. Moreover, another recent paper examined the genetic structure of viral populations when rebound occurs rapidly in patients without reservoir-reducing interventions [50], and found diverse lineages in most patients. This finding is consistent with multiple cells exiting the reservoir each day when it is at a baseline size, in contrast to the case of HSCT patients.

At the time of rebound, both patients had viral load measurements taken every 2 weeks with a detection limit of 20 RNA copies/ml. According to the model behind Figure 5, the worst-case scenario for reservoir reseeding by the time virus is detected, assuming rebound begins immediately after the last negative test result, is about 0.5 IUPM for this measurement schedule and the rapid rebound rate seen in these individuals. For Patient A, rebound was detected at only 900 c/ml, suggesting (based on the observed rate of exponential growth) that virus only became detectable around 4 days before and that the reservoir size could have been limited to around 0.05 IUPM if ART had been re-initiated immediately. In reality treatment was not initiated until several days later, when viral load reached around 1.2 x 10^5^ c/ml, and suboptimal adherence led to even higher values before suppression was eventually achieved. In contrast, for Patient B the first positive viral load value was already likely near the pretreatment peak, at 1.9 x 10^6^ c/ml, so even though treatment was initiated promptly leading to rapid viral decline, near-complete reservoir reseeding may have been inevitable.

## Discussion

One of the proposed strategies to cure HIV infection is to reduce or eliminate populations of latent virus that persist despite long-term antiretroviral therapy. The goal of these strategies is to allow individuals to stop cART without experiencing rebound of viremia. However, case studies such as the “Mississippi baby” [8] and the “Boston patients” [7] highlight the difficulties in conducting clinical trials of reservoir-reducing interventions. Most worryingly, these cases demonstrate that latent virus levels can be undetectable with current technology and still cause rebound after long delays. As a result, long-term follow-up during cART interruption is needed to verify viral eradication. In this paper we have used a combination of mathematical and statistical modeling techniques to guide the design and interpretation of clinical trials for novel curative interventions. This framework includes a method to estimate the size of the remaining latent reservoir and the probability that the infection is cured, based on the delay in viral rebound following cART-interruption. The same framework also can be used to determine adequate trial cohort sizes to make these estimates; to optimize the schedule of viral load testing during supervised interruption; and to estimate the extent of reservoir reseeding if rebound goes undetected, which is a risk for any interruption trial.

One of the most useful features of our approach is that it offers a formal way to combine information from pre-interruption assays of the latent reservoir size, even if they are all negative, with observations during treatment interruption. Our analyses predict that even if no infection is detected among approximately 100 million cells assayed, 4 years of follow-up are needed before there is a 50% chance of cure, and over 10 years to be 90% sure. The higher cell samples that could be obtained with leukophoresis (≈ 1 billion) only decrease these follow-up times by around a year. An implicit assumption of this approach is that there is a dynamic equilibrium between latently infected cells in the tissues and those circulating in the peripheral blood. If a reduction observed in the blood does not correspond to the reduction in the tissues, these estimates will be biased. Consistent patient monitoring during treatment interruption is necessary, and how the frequency of sampling should be scaled-back as time progresses and cure becomes more likely is not intuitive. We used the model predictions to design a sampling schedule so that the chance of patient rebound between samples remained at a constant low level. It suggests times after which sampling frequency could be reduced to weekly (5 months), monthly (1.5 years), quarterly (5 years), and annually (10 years). However, even when the risk of rebound between samples is low, if it does occur, the risk of reaching high-level viremia and completely reseeding the reservoir before the infection is detected may be high.

Even when the post-intervention reservoir size is informed by negative assay results and a precise measurement of viral rebound time, there is considerable uncertainty in its size. This uncertainty in residual HIV reservoir size follows from the stochastic nature of viral rebound from small latent reservoirs. This stochastic process produces large variability in viral rebound times, even for patients starting with identical reservoir sizes [10]. As a result, large trial sizes – between 40 and 150 individuals – will probably be necessary to reliably estimate the reservoir-reducing potential of a new therapy and to compare this across interventions.

Many of our estimates (Figure 2-4, Table 1: “Predicted from model”) rely on a mechanistic model of rebound dynamics that was developed and detailed in a previous manuscript [10]. The model we developed treats the latent reservoir as a homogeneous population of cells, in which activation and subsequent release of infectious virus particles is assumed to be a probabilistic event occurring independently for each cell. While sufficient data does not yet exist to validate this model formally, independent confirmation of some of its predictions suggest that the model is informative and relevant. The model independently predicted that rebound may occur after months to years after cART cessation as observed in the Mississippi and Boston cases [7, 8]. Genetic analysis of rebounding viral lineages suggests that rebound may be caused by multiple lineages when reservoir sizes are high and rebound occurs rapidly [50], but only single lineages when reservoir sizes are undetectable and rebound is delayed, such as in the Boston patients [7]. These findings are consistent with the model, in which delays in rebound were due to a decrease in the daily reactivation rate of latently infected cells.

Quantitative predictions of the model depend on parameters governing rebound dynamics. As described in the first section of Results, four parameters play an important role: the half-life of the pool of latently infected cells, the activation rate of latently infected cells, the viral growth rate after activation, and the probability that a single activated cell establishes a growing infection. Biological details of viral and cellular dynamics, including the proliferative mechanisms maintaining the latent reservoir, the length of the “eclipse phase” during which a cell does not produce virus particles, and the number of virus particles produced by a single infected cell, were seen to have only minor effects on rebound [10]. Accordingly, in the current paper, we have not considered these additional details and have confined sensitivity analysis to the four main parameters alone (). There is broad consensus regarding measurement of the half-life of the latent reservoir for patients on ART and the viral growth rate following rebound: Longitudinal studies of latent reservoir size show a half-life of about 44 months for the typical patient [15, 16], and viral load dynamics show an exponential growth rate of about 0.4 per day for the typical rebound event [17, 18]. Direct estimates of the latent cell activation rate and the establishment probability are difficult to make, and there is ongoing debate regarding methods to measure them [51, 52]. The problem is essentially one of observing extremely small numbers of cells *in vivo*: in an ART patient with a fully-suppressed viral infection, there may be fewer than 100 actively infected cells present at any one point in time [10], and it is not feasible to count, much less track the fate of, each one of these cells. For now, the field must pursue indirect measurements [10, 24], informed by viral dynamics [17, 18], population genetics [21–23], in *vitro* virology [19, 20, 53], and the timing of viral rebound [51, 52]. Improvements in these estimates will be invaluable to the field, since indirect, model-based methods for interpreting treatment interruption trials will be necessary until ultrasensitive assays for the latent reservoir are created.

The statistical framework presented here could be adapted to any mathematical model which similarly forecasts rebound. In particular, it can be applied to the model recently proposed by Pinkevych et al. [51], who argued that the product of two of the parameters described above – the activation rate of latent cells and the probability that a single activated cell manages to establish a growing infection – is much lower than we have suggested (0.17 cells per day, versus 4 cells per day). This lower estimate implies that several months’ delay in rebound could be achieved with less than one log-reduction. For reasons fully detailed in a separate publication [52], we believe that the Pinkevych et al. model underestimates this product by not appropriately accounting for interpatient variation in latent reservoir size and viral growth rates when interpreting data from treatment interruption studies. Our analysis therefore employs our original model and parameter estimates.

An additional limitation of the model is that it assumes that the intervention reducing reservoir size is homogeneous and does not change the parameters governing reservoir or virus dynamics. While the latter assumption seems reasonable for treatment with some latency-reversing drugs, HSCT drastically perturbs the immune system, which may in turn alter the rate at which latently infected cells proliferate, die, or reactivate. It may also alter the growth patterns of viral lineages that stem from reactivated cells. For our analysis of reservoir size and rebound times, we assumed no change in these rates. The one parameter we could easily measure – the viral growth rate during rebound – did appear to be larger than the value in non-transplanted individuals, perhaps due to a loss of HIV-specific immunity. This parameter, however, has a small effect on the time to rebound relative to others [10]. If there were evidence to suggest the reactivation rate of latent cells was higher in these patients due to GVHD or other effects, then the observed rebound times would be consistent with lower reservoir sizes than estimated in our analyses. Another consequence of the simple way in which the model treats the immune system is that it cannot predict rare phenomena such as post-treatment control [54, 55]. This shortcoming, however, may have little effect on prediction of rebound in the large majority of patients. We have assumed that the latent reservoir is a homogeneous population of cells, because at this stage little information is available about phenotypically different compartments and how these may vary in response to LRAs. However, one alternate scenario is easy to study with our model. If the reservoir consists of two compartments – one which is completely reactivated and cleared by the LRA, and one which is not affected at all –then the results our identical to the ones presented here, as long as the reduction efficacy is interpreted as the fraction of latent cells that reside in the unaffected compartment.

We used a separate model, not relying on the above assumptions, to estimate the extent of reservoir reseeding during rebound (Figure 5) and the number of donor cells infected by ongoing replication during engraftment (Table 1). This model predicts the number of latently infected cells using longitudinal viral load and CD4 count data, based on regression coefficients estimated from a cohort observed during acute infection [26]. Although this model is mechanistically grounded in viral dynamics theory [56] and empirically grounded by cohort data, its predictive ability is imperfect. The best-fit model from Archin et al [26] was only able to estimate reservoir sizes to within approximately 1-log of the observed values, and we have used this finding to put error bounds on our estimates. Additionally, the log-linear regression Archin et al used to estimate latent infection rates deviates from the linear relationship predicted by viral dynamics theory, suggesting that the theory behind these predictions is incomplete and that extrapolating beyond the range of observed data, as we’ve done here, may be problematic.

Given current data and understanding of HIV infection, we have the barest estimate of the number of donor cells that became infected (despite administration of cART) and entered latency during HSCT in the Boston patients (Table 1). The existence of ongoing viral replication during cART is a hotly debated topic, but few studies have attempted to place an upper bound on the rate at which new infection could occur [41]. Observations that viral evolution is limited during treatment [35, 36] and that treatment intensification fails to reduce viral load [37, 38] suggest a very low infection rate. Additionally, the long delays in rebound observed in both Boston patients suggest that any ongoing replication must have been transient, since replication at the time of cART-interruption would have resulted in rapid rebound.

Progress in HIV cure research is challenged by major uncertainties — uncertainty over the size and persistence of the infected cell population that must be eliminated to achieve a cure, uncertainty over the immediate effect of potential curative interventions, and uncertainty over the long-term benefits and risks of such interventions. A mathematical framework that incorporates mechanistic modeling and Bayesian statistics will play a key role in managing these uncertainties, enabling the design and interpretation of clinical trials.

## Methods

### Ethics Statement

This manuscript involves the analysis of previously published data from human studies, and details of the ethical approval can be found in the original publications [6, 7].

### Bayesian approach to interpreting trial outcomes

If *q* is the reduction efficacy, defined as the fraction of latent cells remaining after reservoir-reducing therapy, then we define *S*(*t* | *q*) be the fraction of patients who have not yet rebounded at a time t after treatment is stopped. Since we do not know the true function *S*(*t* | *q*), we rely on the predictions calculated from the model. For any fixed q value, the function *S*(*t* | *q*) describes “survival curves” that can be calculated from the model (Figure 1C). The details of the model dynamics and parameter estimation are described in detail in a previous manuscript [10].

We first approach the question of how to estimate the amount by which the reservoir has been reduced, when we cannot measure reservoir size directly from assays but have observed delayed viral rebound. We consider a patient who has still not experienced viral rebound after *τ* days after ART-interruption (rebound time *t_r_* is greater than *τ*). We define *P(q)* to be the probability distribution for our existing knowledge about the likelihood of a given *q* value being the true value (the *prior*). This function can be estimated using data from experimental assays, as detailed in the next section. The function *S*(*τ* | *q*) forms the *likelihood* of the Bayesian approach, which gives the probability of the data (*τ*) given a particular model (the unknown reduction-efficacy parameter *q*). Then, the *posterior probability distribution* for the reduction efficacy *q* becomes:

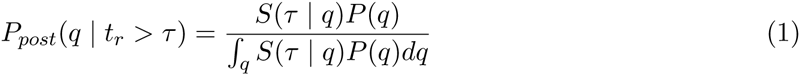

Alternatively, if a patient rebounded *at* a time *τ*, then the formula is slightly modified. The likelihood function for the time of rebound is given by the slope of the survival curves, *∂S*(*τ* | *q*)/*∂t*, so the posterior probability for the reduction efficacy *q* becomes

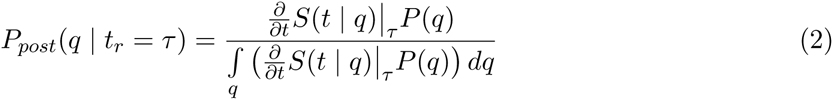

We next turn to determining the probability that a patient is cured (i.e. will never experience viral rebound, even if they were to live forever), as a function of the current time they have been off cART without rebounding. Equation (1) gives the posterior probability distribution for the reduction efficacy *q* when a patient has survived without rebound for a time *τ*, for a given prior distribution. We combine this with the fact that the probability of cure is given by the limiting value of the survival curve as time increases to infinity, *p_cure_(q)* = *S*(∞ | *q*), and that the conditional probability of cure, given only those individuals who have not yet rebounded at time *τ*, is *p_cure_*(*q* | *t_r_* > *τ*) = *S*(∞ | *q*)/*S*(*τ* | *q*). Then, we can derive the probability of a cure, given the absence of rebound until time *τ*, as

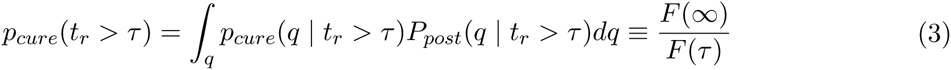

where

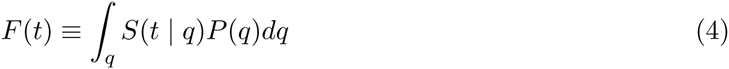

We can also estimate how many patients would be needed in a trial to narrow down the estimated reservoir-reduction to something close to its actual value. We say that this is the trial size required to adequately “power” the study, where we imagine that the main goal of the study is to determine how much some novel intervention reduces the reservoir, and that this can only be done by observing rebound times.

If there are *n* patients in a trial and a set of rebound times {*τ_i_*} = {*τ*_1_, *τ*_2_… *τ_n_*} are observed, then the posterior distribution for the reduction efficacy *q* is given by:

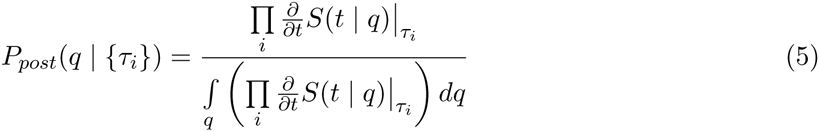

Here we assume a uniform prior, as a worst case scenario where no other information on reservoir size is available. If *q* is high enough so that after some time *τ_max_, m* patients still have not rebounded, then the estimated reduction efficacy becomes

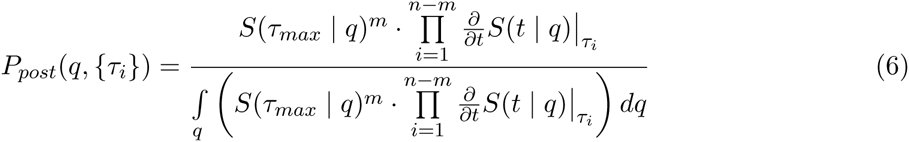

The required number of patients *n* can be calculated for particular requirements on the posterior estimate, by repeatedly simulating patient data and using it to construct *P_post_.* For example, we defined a study as adequately powered if 95% of trials of this size would result in 95% (centered) credible intervals for *q* that that contain the true value and are less than 1 log wide. We determined *n* by doing a binary search though trial sizes from the list {5, 10, 15, 20, 25, 20, 40, 50, 60, 70, 80, 90, 100, 150, 200, 250, 300, 400, 500, 600, 700, 800, 900, 1000}. Here we assume *τ_max_* = 10 years.

We additionally use these methods to estimate the optimal frequency of sampling subjects in a cART-interruption trial. One method to optimally choose sampling time points is to determine intervals between which the chance of a participant experiencing viral rebound is equal.

If the reduction efficacy *q* is known, then the probability that someone who has survived until timepoint *t_i_* rebounds before time *t_i+1_* is

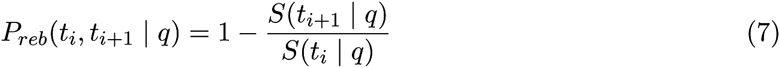

However, in the general case that *q* is unknown, we must again weight this probability with the posterior likelihood of *q*, given that the patient has already survived without rebound until time *t_i_*

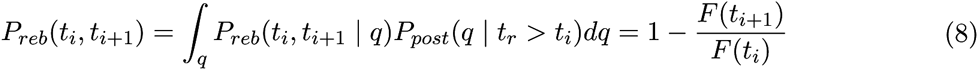

Where *F*(*t*) is defined as in (4). We set *P_reb_*(*t_i_*, *t*_*i*+1_) to the pre-defined “tolerance” δ - the fraction of suppressed trial participants expected to rebound between each sampling timepoint. Using t_0_ = 0 and *F(t_0_)* = 1, we can iteratively solve for *t_n_* using the implicit equation *F(t_n_)* = (1 – δ)^*n*^.

We do not have analytic functions for the model output *S*(*t* | *q*), which means we cannot analytically calculate any of the integrals described above. Instead we use the simulated model output to numerically construct *S*(*t* | *q*), using 10,000 patients for each *q* value and *q* values between 10^−6^ and 1 in increments of 0.05 logs. We then calculated the integrals as Reimann sums.

### Deriving priors

The Bayesian approach to estimate the post-treatment reservoir size (*q*) as a function of rebound time can incorporate any prior knowledge we may have about the likely reservoir size (*P(q)*). This prior distribution can be constructed based on the results of virologic assays that were used to measure the size of the reservoir before cART was interrupted.

Briefly, to construct *P(q)*, we imagine an assay was performed that measured the frequency of latently infected cells in a sample (e.g. by viral outgrowth or PCR). We assume that cells used in the assay were randomly sampled from a much larger pool in a patient’s body. We can then construct the likelihood distribution for the true frequency of latently infected cells, based on the actual number of infected cells observed in the sample. We define the true frequency of latently infected cells to be *θq*, where *θ* is the frequency before reservoir-reduction, and *q* is the fractional reduction. If *C* cells are sampled and zero infected cells are observed, then the likelihood for *q* goes as *P*(*q*) ∝ *e^−θCq^*. (This is based on the Poisson approximation to the binomial process when the number of samples is very high and the probability of success is very low). This distribution shows that taking only negative assay results into account, any *q* value is possible, but *q* = 0 (no remaining reservoir) is mostly likely and higher *q* values (larger reservoir sizes) are less likely, although as *q* becomes very small, the distribution flattens out.

More details of this calculation can be found in separate manuscript [14]. Simple browser-based software for calculating the full likelihood distribution for viral outgrowth assays, even when positive results are detected, is available at http://silicianolab.johnshopkins.edu.

Using *C* = 10^8^ and θ = 10^−6^ (i.e. the reservoir was sampled at a level of 100 times the pre-reduction frequency of latency), then if no infected cells are detected, the resulting prior distribution is shown in Figure 2a. Note that on a log-scale, larger reductions (e.g. between 5 and 6 logs) appear less likely, which is simply because the density of numbers between these values is less. The actual individual reservoir size value with the highest likelihood is still *q* = 0 (*Q* → ∞).

Other priors may also be possible, depending on the nature of the assay. The most uninformative prior would be to assume that we have no information on reservoir size before cART interruption, and so every reservoir size is equally likely. We think this is unrealistic, since an interruption trial would likely not be conducted unless the reservoir was undetectable or at least very low by standard assays.

Note that it is common practice in clinical studies to interpret the results of the types of assay described above as a hard cut-off for the reservoir size. It is often said that if no infected cells are detected, then the frequency is less than the inverse of the sample size, i.e. *θq* < 1/*C*, or *q* < 1/*C*θ). For the scenario with *C* = 10^8^ and θ = 10^−6^ used above, this would lead to the conclusion that there was at least a 2-log reduction in the reservoir size. However, this “detection limit” actually represents a very weak confidence interval, because in repeat sampling, it is expected that 63% of all draws will from a sample with an infected frequency at this “limit” will give at least one infected cell.

## Supporting Information

### S1 Text

**Estimating reservoir size with uncertainty in viral dynamics parameters**

### S2 Text

**Estimating reservoir re-seeding during viral rebound**

### S3 Text

**Estimating the size and source of the reservoir post-transplant**

